# Hydrodynamic model of directional ciliary-beat organization in human airways

**DOI:** 10.1101/2020.01.06.896001

**Authors:** Simon Gsell, Etienne Loiseau, Umberto D’Ortona, Annie Viallat, Julien Favier

**Affiliations:** Aix Marseille Univ, CNRS, Centrale Marseille, M2P2, Marseille, France; Aix Marseille Univ, CNRS, CINAM, Marseille, France

## Abstract

In the lung, the airway surface is protected by mucus, whose transport and evacuation is ensured through active ciliary beating. The mechanisms governing the long-range directional organization of ciliary beats, required for effective mucus transport, are much debated. Here, we experimentally show on human bronchial epithelium reconstituted *in-vitro* that the dynamics of ciliary-beat orientation is closely connected to hydrodynamic effects. To examine the fundamental mechanisms of this self-organization process, we build a two-dimensional model in which the hydrodynamic coupling between cilia is provided by a streamwise-alignment rule governing the local orientation of the ciliary forcing. The model reproduces the emergence of the mucus swirls observed in the experiments. The predicted swirl sizes, which scale with the ciliary density and mucus viscosity, are in agreement with *in-vitro* measurements. A transition from the swirly regime to a long-range unidirectional mucus flow allowing effective clearance occurs at high ciliary density and high mucus viscosity. In the latter case, the mucus flow tends to spontaneously align with the bronchus axis due to hydrodynamic effects.

## Introduction

Billions of microscopic active cilia on the surface of the bronchial epithelium propel a viscous fluid, called mucus. Its function is to capture and eliminate inhaled pathogens, allergens and pollutants^1^. This process, referred to as mucociliary clearance, is impaired in respiratory diseases such as primary dyskinesia, severe asthma^2^, COPD^3^ and cystic fibrosis^4,5^, which affect hundreds of millions of people worldwide.

The transport of mucus requires a long-range coordination of ciliary beat directions along the airways. In *in-vitro* cell cultures, observations of large-scale circular mucus flows spanning the culture chamber was reported by Matsui et al.^6^. Recently, Khelloufi et al.^7^ showed that this swirly transport is associated with a circular order of the ciliary beating directions underneath the mucus. Furthermore, mucus swirls of various sizes were observed, whose size was shown to scale with the ciliary density. Due to the invasive nature of *in-vivo* experiments, the mechanisms underlying mucus transport and collective dynamics of the ciliary activity remain poorly understood. In particular, the mechanisms involved in the directional self-organization of the ciliary beats over distances spanning the bronchi and trachea are yet to be elucidated. At the tissue scale, the planar polarization of the epithelial ciliated cells, established during development, is expected to determine the rotational polarity of the basal body at the base of the cilia, which sets the direction of beating^8,9^. However, daily variations of ciliary-driven cerebrospinal fluid flow patterns have recently been observed in *in-vivo* mouse brain ventricles^10^. In addition, Guirao et al.^11^ showed that ciliary-beat directions on cell cultures issued from the subventrical zone of newborn mice could be drastically changed by applying an external flow. Finally, the directional collective order of ciliary beats on the multiciliated skin cells of the *Xenopus* embryo could be refined by applying an external flow to skin explants^12^. These phenomena strongly suggest the existence of a coupling between hydrodynamics and long-range ciliary-beat orientation for at least two animal models. These experimental evidences motivate the development of physical models allowing to reproduce such complex ciliary dynamics and to predict the resulting fluid transport. In the context of mucociliary clearance, such models could be highly beneficial for the comprehension, prediction and control of this crucial physiological process.

A variety of physical models have been developed to describe collective behaviors in groups of interacting active individuals, such as insect swarms, bird flocks, fish schools or bacterial colonies^13–16^. These models, deriving from the seminal work of Vicsek et al.^17^, are based on interaction rules between moving individuals, namely avoidance, attraction and alignment. Several works also focused on the self-organization of cilium-like active structures attached to a substrate, specifically addressing their collective temporal synchronization and the possible emergence of waves^18,19^. Guirao and Joanny^20^ proposed a theoretical model of directional ciliary organization based on the interaction between isolated forces (stokeslets) uniformly distributed over a substrate. The model predicts alignment of the forces, but it does not account for the emergence of circular patterns as observed by Khelloufi et al.^7^. Moreover, this type of modeling assumes small ciliary densities and linear flow behavior, and it does not allow extension of the hydrodynamic model to complex rheologies. An alternative approach, based on the Navier-Stokes equations, is proposed in this work.

Here, we experimentally show on human bronchial epithelia reconstituted *in-vitro* that, in presence of a ciliary-driven mucus flow on the epithelial surface, directions of ciliary beats tend to align progressively along mucus streamlines. Conversely, upon mucus removal, the orientational order of ciliary beat is lost. Based on these observations, we propose a two-dimensional hydrodynamic model of ciliary-beat organization where the ciliary orientation is controlled by a streamwise-alignment rule. In this model, the coupling between distant cilia is ensured only by the hydrodynamics, which is computed based on the full flow equations. The model relies on two independent physical quantities, namely (i) the ciliary density and (ii) the viscosity ratio between the mucus and the periciliary layer. The simulations predict the emergence of mucus swirls whose typical size is consistent with experimental measurements. At large ciliary density and mucus viscosity, a transition to long-range unidirectional mucus flow occurs. In this latter regime, we show that in a virtual bronchus the direction of transport aligns with the distal-proximal axis.

## Results

### Ciliary beat reorientation in human bronchial epithelium cultures

The experimental system consists of well-differentiated human airway cultures at the air-liquid interface (ALI). These cultures exhibit the pseudo-stratified structure typical of bronchial epithelium, with basal cells, goblet cells that produce mucus, and ciliated cells with beating cilia^7^. As described by Khelloufi et al.^7^, the ciliogenesis is accompanied by the emergence of swirly mucus flows. The ciliary density on mature epithelium reaches around 70% of the epithelial surface. At high ciliary densities, the mucus is transported over macroscopic distances (see movie 1) thanks to the continuous beating of cilia. We image both the flow of mucus and the beating cilia underneath (movies S1 & S2). We then quantify the mucus flow direction as well as the beating directions of cilia (Fig. 1 and movie 2). Locally the cilia does not necessarily beat along the mucus flow direction (Fig. 1(a)) but they present an average direction at large scale. Yet, we show that the ciliary beating directions tend to align along the direction of mucus flow (Figs. 1(a,b) and movies S1 & S2) in a slow process of a maximum 35-40° angle reorientation within 24 hours (see angle distributions in Figs. 1(d,e)). In contrast, when mucus is washed out from the surface of the cell culture and replaced with culture medium, we observe within three days a loss of the coordination of beating directions (Fig. 1(c)), highlighted by the emergence of numerous discontinuities in the director field of the beating cilia. At this stage, only small patches of cilia attached to neighboring ciliated cells present the same beating direction. The characteristic diameter of these patches is around 10 to 20 μm, corresponding to groups of 5 to 15 cells. These patches can be seen as unit elements of beating direction, likely constituted by ciliated cells having a common cell polarity. There is no transport of fluid at large scale anymore. These observations disclose the existence of a strong mucus-cilia interplay and that mucus flow triggers an active response of cilia driving their reorientation.

**Figure 1.**
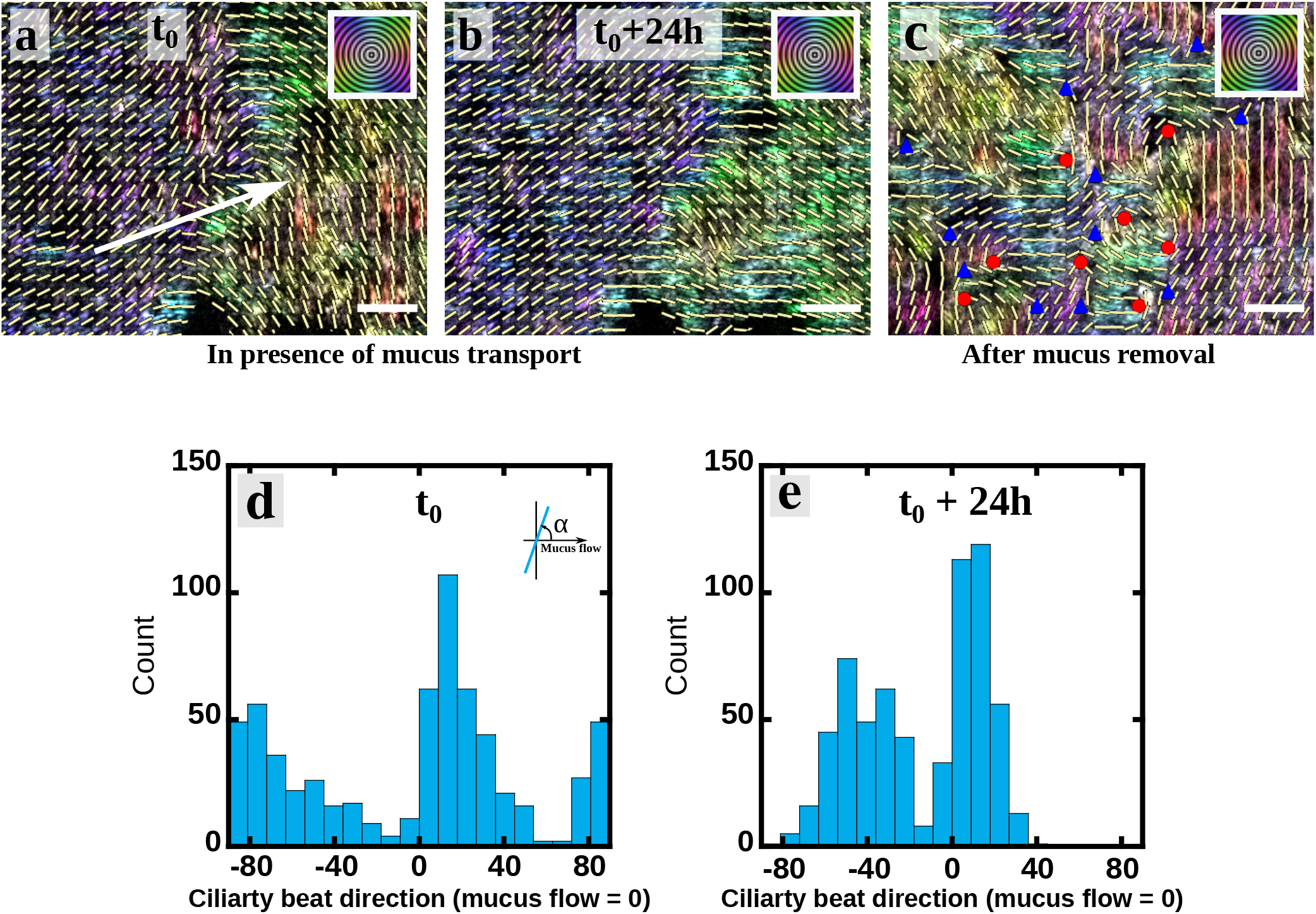
Mucus transport reorients the beating direction of cilia. (a-b) Orientation maps of beating cilia at 24h interval while some mucus is transported above. The colors code for the direction of beating according to the orientation scale on the top right hand corner. The director field of the beating direction is plotted as an overlay and is averaged in boxes of 5 μm x 5 μm (~ size of a ciliated cell). The white arrow indicates the averaged direction of the mucus transport in the considered field and is determined from movie 1. Within 24 hours some reorientations occur, in particular on the right-hand side of the image, and the beating direction can rotate up to 30-35^°^. (c) Orientation map of beating cilia 3 days after washing out the mucus with culture medium. The field of view is the same than in a & b. Colored triangles and circles indicate discontinuities in the director field. Scale bars are 20 μm. The three field of views are representative of the whole culture chamber. The movie of the beating cilia corresponding to the three panels is movie 2. (d,e) Angle distributions of ciliary beats corresponding to the figures (a) & (b) respectively. The 0^°^ orientation corresponds to the mucus flow direction. In presence of a transported mucus above the cilia, we observe a dynamic reorientation of the ciliary beat directions. Over 24h the two peaks of the distribution get more narrow and there is a shift of the two peaks toward the direction of the mucus flow measured from movie 1.

### Hydrodynamic model of ciliary-beat organization

We assume that the hydrodynamic forces generated by the mucus flow drives the observed long range self-organization of the directions of ciliary beats, and propose the following two-dimensional model to examine this hypothesis. It is schematized in Fig. 2(a).

**Figure 2.**
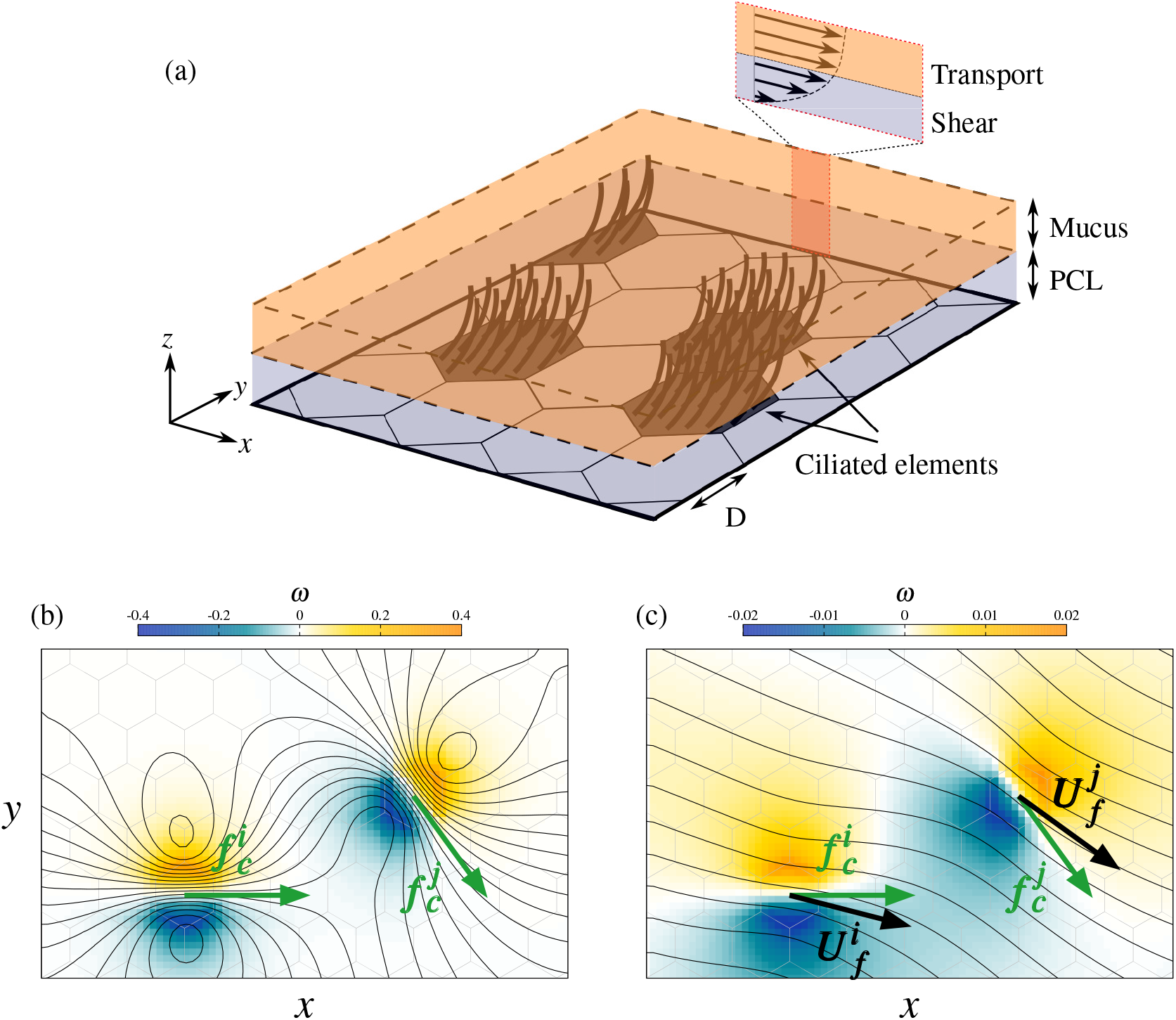
Model of ciliary-driven mucus flow. (a) Schematic view of the model of bronchial epithelium. (b,c) Typical flow in the vicinity of two ciliated elements *i* and *j*, for (b) *λ =* 1 and (c) *λ* = 5. The flow is visualized by streamlines and iso-contours of the flow vorticity. The hexagonal mesh is represented by gray lines. The ciliary forces 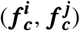 are indicated by green arrows. In (b), black arrows show the locally-averaged flow velocities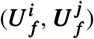.

The main principles of the model are the following. Locally, ciliary beats exert a force on the mucus. Considering a simple modeling of this force, the overall mucus flow resulting from the joint forcing of all ciliated cells is computed by solving the Navier-Stokes equations. The model is based on a streamwise-orientation hypothesis: ciliary beats, and the associated local forces, reorient in response to the computed mucus flow and gradually align with the local flow direction. Such system, initialized with random orientations of the ciliary beats, thus evolves towards a self-organized solution. Distant cilia interact with each other through their individual contribution to the overall forcing field driving the global mucus flow.

In practice, the plane epithelium surface is discretized using hexagonal unit elements, which represent cells or groups of cells (see Fig. 2(a)). Their side length is denoted by *D.* Two types of unit elements are considered, namely multi-ciliated cells, or groups of neighboring multi-ciliated cells that exhibit a joint ciliary-beat direction, and *passive* elements that represent club and goblet cells and, possibly, dysfunctional ciliated cells. We chose a random spatial distribution of these elements on the epithelium, in agreement with observations^11^.

The surface fluid consists of two layers, a low-viscosity periciliary layer (PCL), in which cilia beat, and a high-viscosity mucus layer^1^. The PCL thickness is about the cilia length. The PCL is a shear region; the fluid velocity, in the plane parallel to the epithelium, is zero on the epithelial surface and is equal to the mucus velocity at the PCL/mucus interface^21,22^(see Fig. 2(a)). The mucus layer is modeled as a transport zone. Its velocity is mainly parallel to the epithelial plane and is uniform along the *z* axis, as schematized in Fig. 2(a). Overall, the dynamics of the mucus layer is modeled as a two-dimensional flow developing over a slip layer. This flow is driven by local propelling forces arising from ciliary beating. The forcing is spatially averaged over one or several neighboring multi-ciliated cells (unit element), and temporally over a time scale larger than the ciliary beating period.

Mucus is modeled as a Newtonian fluid, characterized by its density *ρ_m_* and dynamic viscosity *μ_m_.* The flow is governed by the two-dimensional incompressible Navier-Stokes equations. The flow momentum writes

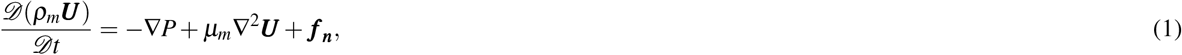

where 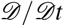 denotes the material derivative, and ***U*** and *P* are the flow velocity and pressure. The body force ***f_n_*** is the momentum source/sink modeling the interactions with the epithelial surface. Two types of momentum sources are considered. The first one results from the shear stress in the PCL; it linearly depends on the mucus velocity ***U***, namely ***f_v_*** = −*κ**U***, with *κ* a parameter called the PCL friction coefficient. The second force component, called the ciliary force and denoted by ***f_c_***, results from the ciliary beating; it is non-zero on ciliated elements, and vanishes elsewhere. Its magnitude |***f_c_***| is the same on all ciliated elements and is considered to be time independent, as the ciliary forcing is averaged over several beating periods. Overall, the total momentum source in Eq. (1) is expressed as ***f_n_ = f_v_ + f_c_***.

The orientation of ***f_c_***, defined by the angle 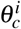 on the *i*-th ciliated cell, is driven by a streamwise-alignment rule. The angle difference between the local flow and the beating direction is 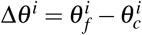, where 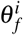 is the flow velocity averaged over the *i*-th ciliated element. The alignment rule follows

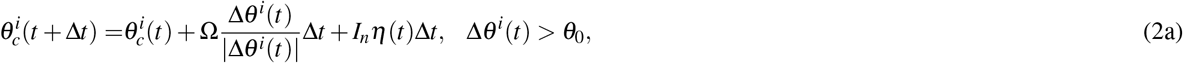

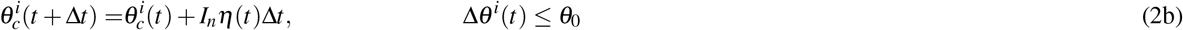

where Δ*t* is the model time step, Ω is a fixed angular velocity, *θ*_0_ is an angle threshold, *I_n_* is the noise intensity characterizing rotational diffusion and *η* is a normalized Gaussian noise. The angle *θ*_0_, which is kept to very small values in this study, is introduced in order to allow the emergence of steady solutions 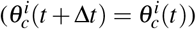 in the absence of noise. This alignment rule provides the hydrodynamic interaction mechanism between ciliated elements. More details on the implemented alignment rule are reported in *Methods*.

Solutions of Eq. (1) are found using numerical simulations (see *Methods*). Periodic conditions are set at the boundaries of the computational domain. The ciliated elements are randomly placed and their total number is set according to the prescribed ciliary density 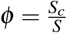, where *S_c_* designates the ciliated area and *S* is the total area. All computations are initialized with random orientations of the ciliated elements.

### Numerical and physical parameters

As noise is not expected to be a dominant mechanism in the real system, the focus is first placed on the model behavior when *I_n_* = 0. In this case, Eq. (2) can evolve to a steady solution where all ciliary elements are close to aligned with the local flow, as defined by Eq. (2)b. These steady organizations, once reached, do not depend on the angular velocity Ω. The latter parameter can thus be considered as a numerical parameter. The threshold angle *θ*_0_ is kept to a small value, so that its influence on model solutions can be neglected.

The PCL friction coefficient *κ* is assumed to relate to PCL properties. From dimensional analysis, it writes 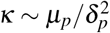, where *μ_p_* and *δ_p_* are the dynamic viscosity and thickness of the periciliary layer. In this model, mucus transport results from the competition between the ciliary forcing and the friction force. Accordingly, a typical mucus flow velocity can be derived from |***f_c_***| and *κ*, namely *U*_0_ = |***f_c_***|/*κ*. This indicates that the mucus velocity is assumed to mostly depend on the ciliary forcing and PCL properties.

Overall, the model mainly relies on six physical parameters, namely *ρ_m_, μ_m_, ϕ, D*, |***f_c_***| and *κ*. Following the Buckingham theorem, the number of parameters can be reduced to three non-dimensional and independent physical parameters, (i) the ciliary density *ϕ*, (ii) the Reynolds number *Re = ρ_m_DU*_0_/*μ_m_* which represents the ratio between inertial and viscous forces and (iii) the non-dimensional interaction length *λ*, defined as

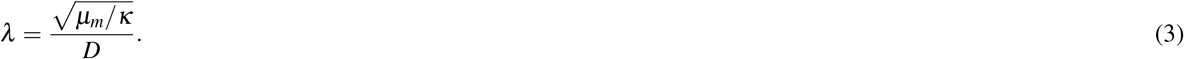

The *λ* parameter represents the typical range of influence of a ciliated element, resulting from the competition between viscous momentum diffusion over the epithelial plane and dissipation due to the friction force ***f_v_***. Indeed, if *λ* is large (i.e. high mucus viscosity), the momentum transferred to the fluid through ciliary beating is rapidly diffused over a large fluid region; the limit case is the solid behavior, where the localized ciliary forcing is instantaneously transferred all over the material. In contrast, the range of influence decreases when *λ* is small, since the flow momentum tends to be damped faster than it is transported over the epithelial plane. Ciliated cells interact with each other only if their separation distance is low enough compared to their individual range of influence. Therefore, *λ* is referred to as hydrodynamic interaction length. Considering the above-mentioned connections between *κ* and the PCL properties, the interaction length is closely related to the mucus/PCL viscosity ratio, namely

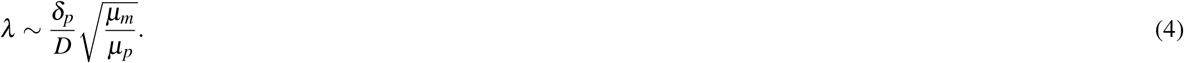

The role of *λ* is illustrated in Figs. 2(b,c), which show the typical flow in the vicinity of two ciliated elements, denoted by *i* and *j*. The non-dimensional flow vorticity is defined as 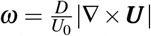. In the figure, high values of the vorticity (either positive or negative) emphasize fluid regions that are sheared along the epithelial plane, as an effect of the ciliary forcing. In this illustrative example, the ciliary force directions are fixed, i.e. the alignment rule Eq. (2) has been ignored. For low values of *λ* (Fig. 2(b)), shear regions remain localized in the vicinity of the ciliated cells, and streamlines crossing the ciliated elements are mostly parallel to ciliary forces. Therefore, Δ*θ^i^* ≈ Δ*θ^j^* ≈ 0, and Eq. (2b) is satisfied. For larger values of *λ* (Fig. 2(c)), large shear regions are observed, even though of lower magnitude than in the previous case, and flows induced by both ciliated cells tend to interact with each other. Consequently, streamlines are not parallel to ciliary forces, as emphasized by the locally-averaged flow velocities indicated in the figure. In this case, a trigger of the alignment rule Eq. (2) induces a re-alignment of the cells. This illustrates that *λ* is expected to control hydrodynamic interactions between ciliated elements.

To determine a relevant range of *ϕ* and *λ* for the simulations, one has to estimate their physiological range. During the ciliogenesis, the cilia density increases from 0 up to a final value of 70 % which corresponds to the cilia density in the bronchus^23,24^. The relevant values of *λ* can be evaluated from Eq. (4). The size *D* for the unit elements is estimated from the size of the ciliated patches exhibiting a joint beating direction, which, as seen in Fig. 1, is of the order of *D* ≈ 2 × 10^−5^m. The thickness of the PCL has been measured by Button et al.^25^ and is of the order of *δ_p_* ≈ 10^−5^m. It is accepted that the PCL viscosity is close to that of water^25^ *μ_p_* ~ 10^−3^*Pa.s*. The literature reports an important variability of the mucus viscosity. Values ranging from *μ_m_* = 10^−2^*Pa.s* to *μ_m_* = 10^2^*Pa.s* have been reported in prior works^26^. The resulting range of a physiological *λ*, based on (4), is then in the range [1.5,150]. Finally, it should be mentioned that the flow behavior is expected to be governed by viscous effects, i.e. *Re* ≪ 1.

### Three different mucus flow regimes: localized, swirly, long-range aligned

A square domain composed of approximately 10^4^ elements is first considered. Rotational diffusion is neglected (*I_n_* = 0), and the Reynolds number is set to 0.1. The flow solution is computed over ranges of *λ* and *ϕ*, namely *λ* ∈ [1,150] and *ϕ* ∈ [0.1,0.7].

The system dynamics is computed until a steady state is reached, when the ciliary-beat organization and the related mucus flow remain constant in time (see movies S3, S4 and S5 for examples of transient dynamics). In this state, the ciliary forces and the flow are aligned locally. Three regimes associated with distinct flow patterns and ciliary-beat organizations are identified, as illustrated in Fig. 3.

**Figure 3.**
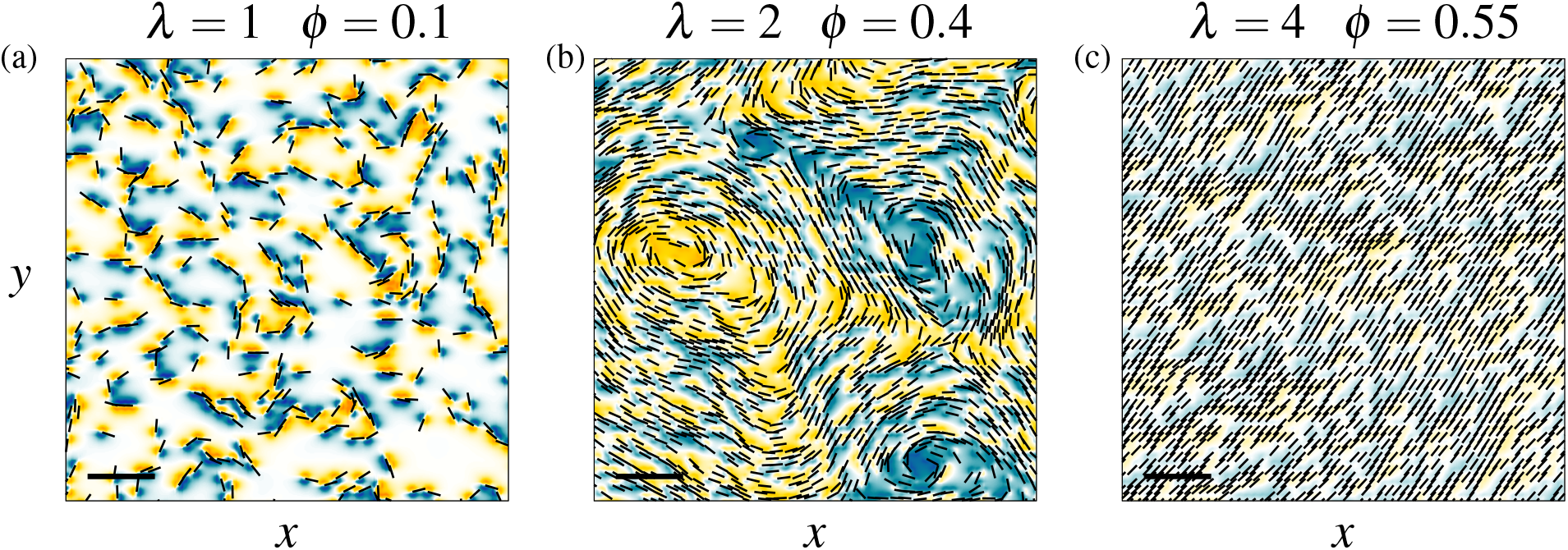
**Visualization of the mucus flow and ciliary-beat patterns** emerging in a periodic square computational domain. Three regimes are identified: (a) a poorly organized regime for *λ* = 1 and *ϕ* = 0.1, (b) a swirly regime for *λ* = 2 and *ϕ* = 0.4, and (c) a fully aligned regime for *λ* = 4 and *ϕ* = 0.55. In each case, the flow is visualized by iso-contours of the non-dimensional vorticity (*ω* = [−0.5,0.5]), and black rods indicate beating directions. Part of the computational domain is shown and the scale bars correspond to 15*D*.

When the ciliary density *ϕ* and the interaction length *λ* are small, typically *ϕ* = 0. 1 and *λ* = 1, a poorly organized regime is observed (Fig. 3(a)). Only local ciliary-beat alignments are observed in regions of high local ciliary densities. No large-scale flow pattern emerges. The flow length scale is of the order of the typical cell size, as indicated by the localized and short-range shear regions close to ciliated elements.

When *ϕ* and *λ* are larger, typically *ϕ* = 0.4 and *λ* = 2 (Fig. 3(b)), a remarkable swirly pattern, associated with local mucus recirculations, emerges. The ciliary beat orientation exhibits circular alignments over length scales much larger than the typical cell size, emphasizing the emergence of large-scale collective behavior. These large vortices are associated with large-scale vorticity regions. Both clockwise and counter-clockwise flow circulations are observed, as shown by the negative (blue) and positive (yellow) values of the vorticity.

When ciliary density and interaction length are further increased (Fig. 3(c)), typically *ϕ* = 0.55 and *λ* = 4, ciliary beat directions tend to fully align over the whole domain. In this case, the flow is unidirectional and almost uniform. As a result, the flow is hardly sheared, as indicated by the low magnitude of the vorticity. The fully aligned regime is the only regime allowing an overall fluid transport. In this case, the average flow velocity can be computed analytically from the momentum balance, expressed as

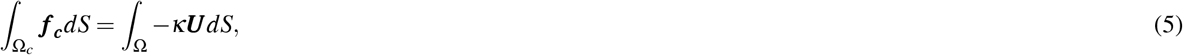

where Ω and Ω_*c*_ are the total and ciliated surfaces. Assuming that the orientation of ***f_c_*** is uniform over the domain, and aligned with the flow velocity ***U***, the average value of the velocity is

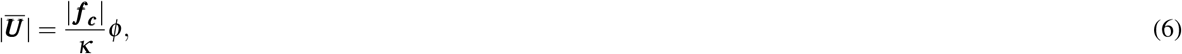

where the symbol ^—^, in this equation and in the following, denotes the space-averaging operator. The mucus velocity is thus expected to linearly increase as a function of the ciliary density. In contrast, the mucus velocity appears to be independent of the mucus viscosity. This is an expected feature of the present model, which involves periodic boundary conditions, thus avoiding any energy dissipation due to boundary layers. Such dissipation is however expected to occur in real systems involving solid boundaries.

In order to quantitatively characterize the three regimes, two physical quantities are defined. The first one is the polarization 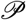,

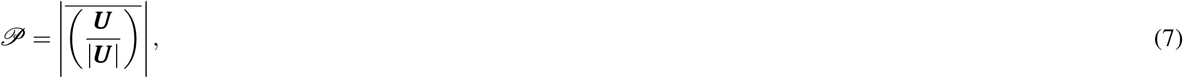

which is a spatial averaging of the unitary velocity vectors and quantifies the orientational order of the overall flow. 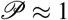 is a signature of a unidirectional flow. The second quantity, Λ, is defined as,

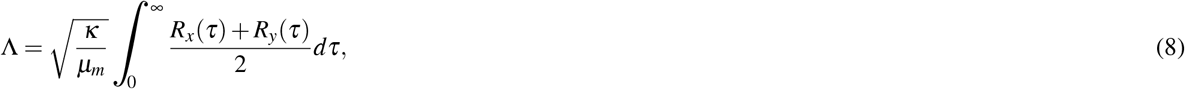

where *R_x_*(τ) and *R_y_*(τ) are auto-correlation functions of the vorticity along the *x* and *y* directions respectively.

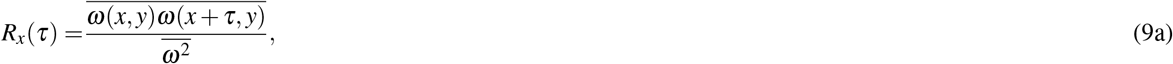

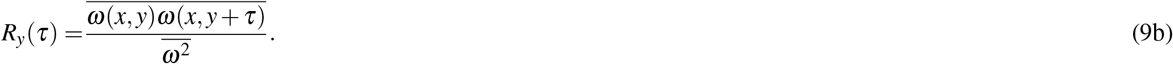

Λ is a non-dimensional integral length. It determines the typical length scale of flow structures. In Eq. (8), it is normalized by 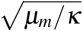, the dimensional interaction length. Therefore, Λ = 1 indicates that the flow length scale corresponds to the range of influence of one ciliated element, i.e. no long-range dynamics occurs. In contrast, Λ > 1 indicates that large-scale vorticity regions are present, as in Fig. 3(b).

The evolutions of 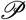 and Λ in the (*ϕ,λ*) plane are plotted in Figs. 4(a) and 4(b), for *λ* ranging from 1 to 5, i.e. the range where regime transitions occur. From these variations, we identify three regions in the (*ϕ, λ*) space: the low-*λ* / low-*ϕ* region corresponds to a poorly organized regime, the low-*λ* / high-*ϕ* region corresponds to a swirly regime, and the high-*λ* / high-*ϕ* region corresponds to a fully aligned regime. We establish the flow regime phase diagram (Fig. 4(c)) by combining the two parameter 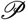 and Λ as following:

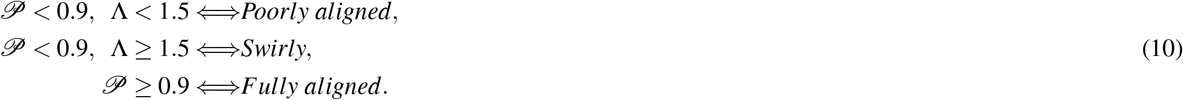

**Figure 4.**
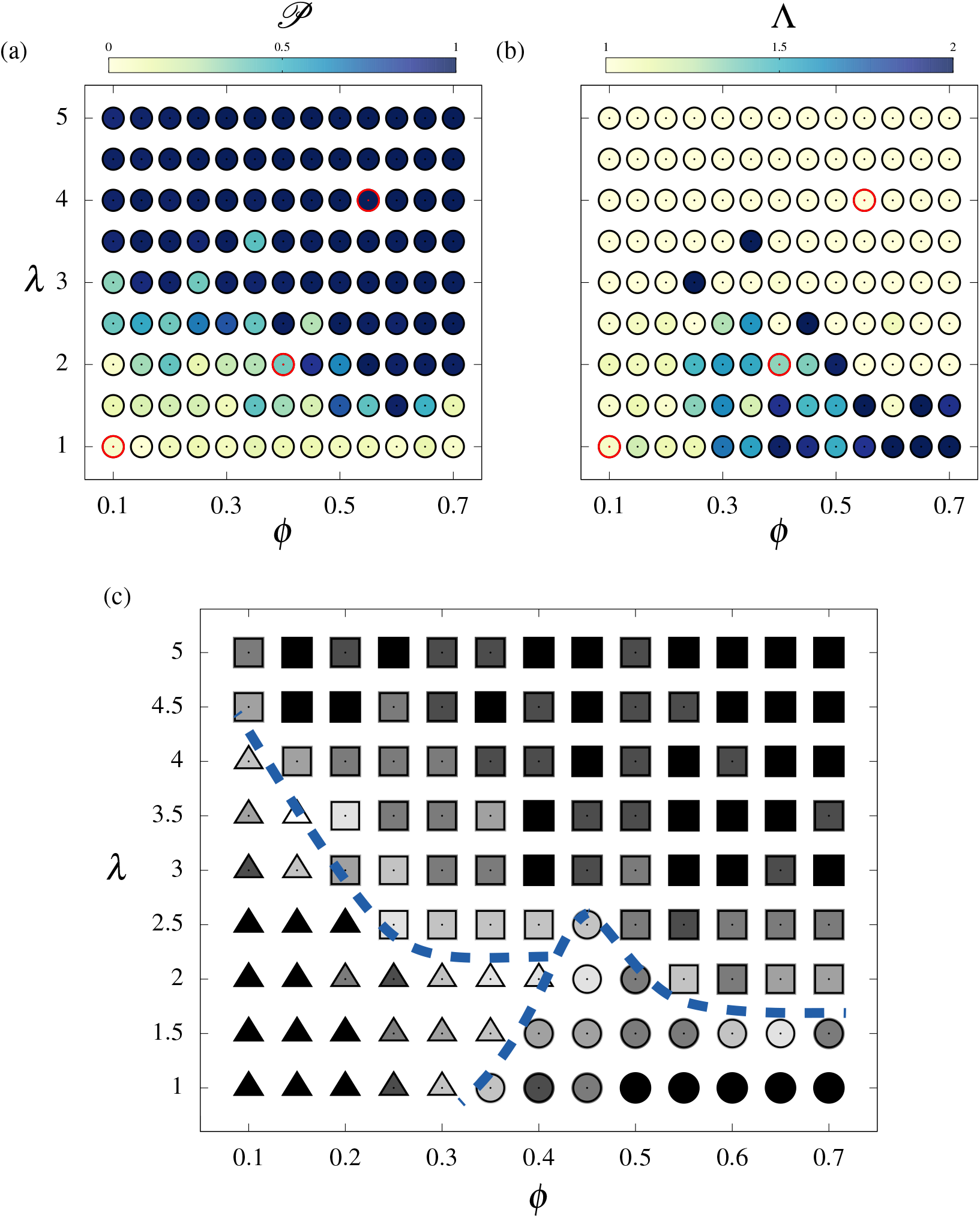
Mapping of the regimes in the (*ϕ, λ*) parameter space. Evolution of the (a) polarization 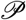 and (b) integral length Λ. The cases plotted in Fig. 3 are indicated by red circles. (c) Overview of the identified regimes based on Eq. (10); at each point, the most frequent regime over a set of ten randomly initialized runs is indicated by a triangle (poorly aligned), a circle (swirly) or a square (fully aligned) symbol. Symbols are colored using the occurrence frequency, ranging from 1/3 (white) to 1 (black). Dashed lines delimitate the three identified regions.

The emerging regimes may vary from one simulation to the other due to the random initialization of the computations. In Fig. 4(c), each point of the phase diagram represents the most frequent regime emerging over a set of ten simulations. The greyscale used in the figure illustrates the sensitivity of the regimes to initial conditions. As expected, flow regimes are sensitive to initial conditions close to the transition regions.

The observed regimes remain robust in the presence of noise. In particular, the swirly regime is almost unaffected by the noise source even when the noise intensity *I_n_* is larger than the alignment velocity Ω (see Eqs. 2), although ciliary disorganization can be achieved for *I_n_* ≪ Ω. Details on the effect of noise on the swirly regime are provided in the *Supplementary information*.

### Swirl size scales with ciliary density and interaction length

The size of the vortices in the swirly regime is determined using the auto-correlation of the flow vorticity, defined by Eq. (9). Additional series of simulations are performed in the region where the swirly regime typically emerges, namely *ϕ* ∈ [0.1,0.7] and *λ* ∈ [1,2.5]. For each point of the (*ϕ, λ*) space, 25 randomly-initialized simulations are carried out. Cases where the number of occurrences of the swirly regime is too small (lower than 10) are discarded. Simulations eventually converging to the swirly regime are selected for the computation of the swirl size. For each case, the average auto-correlation function < *R*(τ) > is computed, and the swirl size is defined as the minimum length scale *τ* satisfying < *R*(τ) >= 0. The swirl size is normalized by *D* and denoted by Λ′.

The evolution of Λ′ as a function of the ciliary density and interaction length is depicted in figure Fig. 5(a). Swirl sizes determined following this procedure are in agreement with qualitative visualizations of the flow. For *λ* = 2 and *ϕ* = 0.4, the measured swirl size Λ′ ≈ 20 is consistent with the swirls depicted in Fig. 3(b). At low ciliary densities, the globally-disorganized solution may include localized swirls, as observed in Fig. 3(a) for *λ* = 1 and *ϕ* = 0.1. The size of these small vortices is consistent with the value Λ′ ≈ 5 plotted in Fig. 5(a). Overall, the swirl size increases as a function of *ϕ* and *λ*.

**Figure 5.**
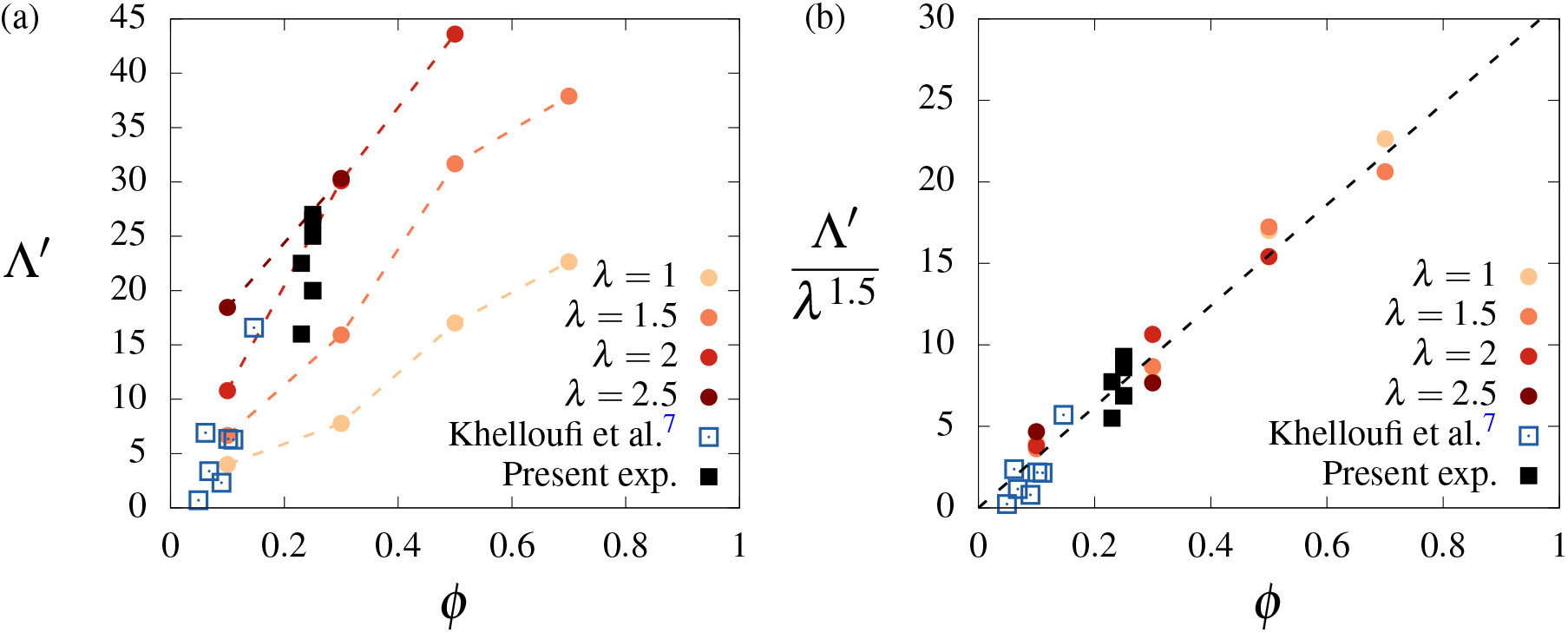
Vortex size scales with *ϕ* and *λ*. The swirl size Λ′ is plotted as a function of the ciliary density for different values of the interaction length in (a), and compared to experimental measurements issued from the present study and from the work of Khelloufi et al.^7^. In (b), all the data collapse on a linear curve when Λ′ is normalized by *λ*^1.5^, illustrating the scaling of Λ′ with *λ* and *ϕ*. Experimental data are normalized using *λ* = 2.

The swirl sizes measured in the present *in vitro* experiments, as well as those reported by Khelloufi et al.^7^, have been included in 5(a) for comparison purpose. Only unconfined results, when the swirls are not impacted by the walls of the culture chamber, are reported here. In the figure, the swirl size has been normalized by *D* = 2 × 10^−5^m, namely the typical size of ciliary patches observed in the experiments (see Fig. 1). Both experimental data sets exhibit a consistent trend that is in agreement with the evolution predicted by the present model, in particular for the case *λ* = 2.

The evolution of Λ′ in Fig. 5(a) follows a trend that is close to linear as a function of *ϕ*, with an increasing slope as a function of *λ*. Assuming a trend Λ′ = *λ^α^ϕ*, a manual fit suggests *α* ≈ 1.5. The evolution Λ′/*λ*^1.5^ as a function of *ϕ* is plotted in Fig. 5(b). A striking collapse of the experimental and numerical data is noted, emphasizing the linear evolution the swirl size as a function of the ciliary density. However, the detailed physical mechanisms involved in these scaling remain to be understood.

### Ciliary beats align with the axis of a virtual airway

To get deeper insight in the role of the domain geometry on the flow direction in the fully aligned regime, we performed a series of 96 randomly initialized computations in a rectangular box with different aspect ratios. The aspect ratio is varied by increasing the domain size in the *x* direction (*L_x_*); the domain size in the *y* direction (*L_y_*) is kept constant. We use a box of the order of 2500 unit elements, with *λ* = 5 and *ϕ* = 0.4 for which long-range fully aligned flow regime is predicted. For a square domain, no preferential flow direction is observed from one computation to the other (Fig. 6(a)). When the aspect ratio of the domain is equal to 2 (Fig. 6(b)), a preferred flow direction emerges along the longest side of the domain. Upon further increase of the aspect ratio *L_x_* = 3*L_y_* (Fig. 6(c)) the effect is even stronger and only a minor part of the independent calculations deviate from the preferential direction.

**Figure 6.**
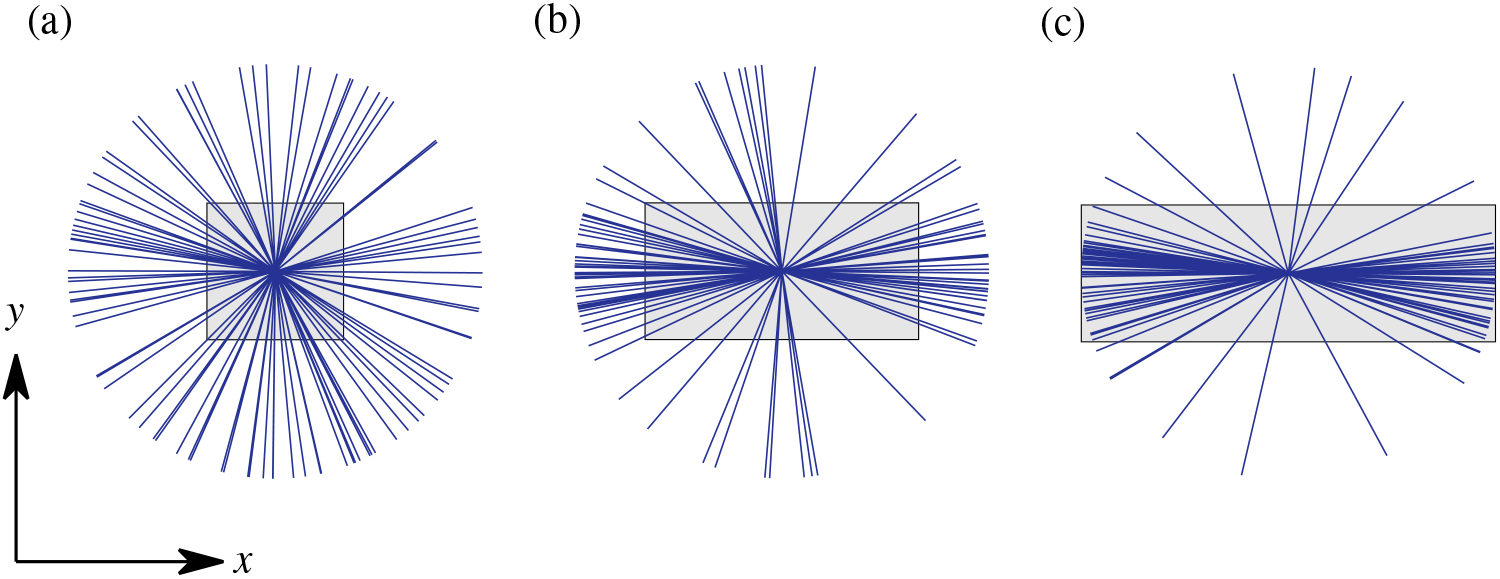
**Anisotropic confinement effect on the fully aligned solutions** in a *L_x_ × L_y_* rectangular periodic domain, for *λ* = 5 and *ϕ* = 0.4: (a) *L_x_ = L_y_*, (b) *L_x_* = 2*L_y_* and (c) *L_x_* = 3*L_y_*. In each case, the space-averaged flow directions issued from a series of 96 randomly initialized computations are represented by blue lines, and a gray rectangle indicates the geometry of the computational domain.

This alignment mechanism relates to an anisotropic confinement effect. Here, the term *confinement* is employed as the size of the periodic domain (*L_y_/D* = 50) has an order of magnitude comparable to the range of influence of a ciliated element, determined by *λ*. Therefore, the periodicity of the domain has an influence on the ciliary organization. When the periodicity is anisotropic, the isotropy of the solution is lost and a preferential direction spontaneously emerges, even though this direction can hardly be predicted *a priori*. The present simulations indicate that the preferred direction is aligned with the longest side of the rectangular domain.

The rectangular geometry is particularly interesting as it is a Cartesian representation of the surface of a circular tube aligned with the *x* axis. The *y* periodicity of the rectangle relates to the azimuthal periodicity of the tube, and thus depends on its diameter. In the *x* direction, the periodic boundary condition ensures that the flow is not constrained in this direction. The possible connections with a natural airway are further discussed hereafter.

## Discussion

We proposed a hydrodynamic model to investigate the emergence of orientational order on a bronchial epithelium. It is based on the description of the two-dimensional mucus flow over an array of self-oriented momentum sources. The orientation of these driving forces is controlled by a streamwise-alignment rule, providing a hydrodynamic coupling between the flow and the ciliary forcing. The model is governed by two independent parameters, the ciliary density and the interaction length, related to the friction over the epithelium and depending on the mucus/PCL viscosity ratio.

The first important result of the model is that it predicts the three main flow regimes experimentally observed: (i) the local flow regime (Fig. 1(c)), (ii) the swirly regime^7^ and (iii) the aligned flow (Figs. 1(a,b)). Experimentally, ciliary disorganization is observed when the mucus is washed using the culture medium, whose mechanical properties are expected to be close to that of water (see Fig. 1(c)). This configuration can be addressed with the present model using *μ_m_ = μ_p_*, leading to *λ* ≈ 0.5. Even at high ciliary densities, only small mucus recirculations are predicted by the model in these conditions (see Fig. S4, which shows the self-organized pattern for *λ* = 0.5 and *ϕ* = 0.7), which is consistent with the disorganization experimentally observed. The long-range swirly regime emerges for intermediate values of *λ*, typically *λ* < 3. The swirl sizes issued from the simulations appear to scale with the ciliary density and interaction length, and show a quantitative agreement with the experimentally measured swirls. The model predicts a fully aligned flow on mature epithelia (*ϕ* ∈ [0.5,0.7]) if *λ* is larger or equal to 2. Physiologically, this minimal value of *λ* is expected to be achieved in human airways since, as reported in the literature and discussed above, the values of *D* ≈ 2 × 10^−5^m, *δ_p_* ≈ 10^−5^m, *μ_p_* ~ 10^−3^Pa.s, and the estimate *μ_m_* ∈ [10^−2^,10^2^]Pa.s lead to the range *λ* ∈ [1.5,150].

Interestingly, the model predicts that the mucus viscosity tends to improve the directional organization of the epithelium, through an increase of the interaction length *λ*, thus enhancing the mucus clearance. On the other hand, mucus viscosity is often expected to have a detrimental effect on the clearance, mainly due to viscous dissipation in highly-sheared regions (boundary layers), that tends to decrease the transport velocity. These antagonistic mechanisms emphasize the possibly complex role of the viscosity in the mucus clearance process.

The role of the ciliary density on long range mucus flow is also quantitatively described by the model. It could therefore be of interest for diseases affecting ciliary motility, as primary dyskenesia. Such a model can open the way to the prediction of the minimal density and spatial distribution of active cilia required to ensure a long-range mucus transport, i.e. the number of ciliated cells that has to be repaired on the airway.

A remarkable result of the model concerns the flow in rectangular domains with periodic boundary conditions. As stated above, this geometry can represent the 2D projection of a circular tube, similar to a natural airway. This comparison with a virtual human bronchus can be further analyzed by considering typical geometrical quantities. Indeed, an effect of the azimuthal periodicity on the directional organization is expected to occur only if the periodicity length and the interaction length have comparable orders of magnitude. Based on the morphometric model of Weibel et al.^27^, the typical cross-section perimeter of human airways is expected to range from *L* ≈ 10^−4^m to *L* ≈ 10^−2^m^28^. By using *D* ≈ 2 × 10^−5^m, the non-dimensional periodicity length is therefore expected to range from *L/D* ≈ 5 to *L/D* ≈ 500. This range is of the same order of magnitude as the expected range of interaction lengths in natural airway, namely *λ* ∈ [1.5,150] as discussed above, and interaction between both length scales is thus possible. The results of the model therefore suggest that, in addition to the biologically driven phenomena occurring during airway development, the distal-proximal alignment of ciliary beats is favored by hydrodynamic interactions in a tube geometry.

## Methods

### Experimental methods

#### In-vitro bronchial epithelium

Mature commercial cultures (MucilAir) of reconstituted human bronchial epithelium from primary cells, were bought from Epithelix (Switzerland). The tissue is cultured at the Air Liquid Interface in a transwell of 6 mm in diameter. The apical side of the tissue is at the air interface and the basal side is in contact with the culture medium (Epithelix MucilAir culture medium) via a porous membrane. The culture medium (700 μl) is replaced every two or three days. For experiments in absence of mucus, we removed the mucus by performing an apical washing. We added 200 μl of culture medium on the apical side for 10 minutes, followed by three rinses with the culture medium. After an apical washing, a thin layer of medium culture (~ 10 − 40 μm) remains on top of the cilia. Experiments were repeated on three different cultures.

#### Imaging

We imaged the cultures in bright field on an inverted Nikon Eclipse Ti microscope with a x20 objective and a Luminera infinity camera. The temperature is maintained at 37 ^°^C and a humidified air flow of 5% CO2 is applied to maintain a physiological pH. To image the culture at the same place on different days, we used a motorized stage calibrated in the xy plane and saved the different positions.

#### Image processing

Beating directions of cilia were quantified using an in-house image processing routine applied on videomicroscopy acquisitions (40 fps). We first performed a standard deviation projection over 40 frames to reveal the trajectory of the tip of the cilia. Then we implemented in python the algorithm described in Ref.^29^ based on the structure tensor computation.

### Numerical methods

The mucus flow, governed by Eq. (1), is simulated using a *D*2*Q*9 Bhatnagar-Gross-Krook lattice-Boltzmann method, which is second-order accurate in time and space^30^. The effect of the body force ***f_n_*** is ensured by the external forcing scheme proposed by Guo et al.^31^. The physical domain is discretized on a uniform and Cartesian numerical grid. The distance between two neighboring nodes, in the *x* or *y* direction, is denoted by Δ*n*; it is set to Δ*n* = *D*/5, where *D* denotes the side length of the hexagonal elements (see Fig. S1). An hexagonal element thus typically contains 65 fluid nodes. At each time step, the flow velocity is averaged in each ciliated elements, and the angle of the ciliary forcing is updated following Eq. (2). The parameter **Ω**, which drives the transient re-orientation of the ciliated elements but does not affect the final steady solution in the absence of noise, is set to **Ω** = *U*_0_/*D*, unless otherwise stated. It is recalled that *U*_0_ designates the typical mucus velocity. A small threshold angle is employed, namely *θ*_0_ = 2ΩΔ*t* = 0.004 radians using the present numerical parameters, so that its influence on model solutions can be neglected. The Gaussian noise component *η* is updated at each time step using a Box-Muller transform. More details on the numerical methods are provided in the *Supplementary information*.

## Acknowledgements

The project MACBION leading to this publication has received funding from Excellence Initiative of Aix-Marseille University - A*MIDEX, a French Investissements d’Avenir programme, the LABEX MEC, and the SINUMER project (ANR-18-CE45- 0009-01) of the French National Research Agency (ANR).

## Author contributions statement

S.G., E.L., U.D., A.V. and J.F. designed research; S.G. developped the model; E.L. performed the experiments; S.G., E.L., U.D., A.V. and J.F. analyzed the data and wrote the paper.

## Additional information

**Supplementary information** accompanies this paper.

**Competing interests:** The authors declare no competing interests.

