## Supplementary information for "Hydrodynamic model of directional ciliary-beat organization in human airways"

**Movie S1:** Transport of mucus above beating cilia corresponding to the field of view of the figure 1(a) & 1(b). The movie is accelerated 150 times. The still image on the right panel corresponds to the projection of 20 frames of the movie of mucus transport using a standard deviation projection method to determine the streamlines of the mucus. The scale bars are 20 m.

**Movie S2:** Beating cilia on in-vitro reconstituted human bronchial epithelium cultures. Left, center and right panels corresponds to the field of views of the figure 1a, 1b and 1c respectively. The total time of the movie is 2.5s and scale bars are 20 m.

**Movie S3:** Time-evolution of the mucus flow and ciliary-forcing directions, for $\phi = 0.5$ and $\lambda = 1$. The flow is visualized using iso-contours of the flow velocity magnitude, and black rods indicate beating directions.

**Movie S4:** Time-evolution of the mucus flow and ciliary-forcing directions, for $\phi = 0.5$ and $\lambda = 2$. The flow is visualized using iso-contours of the flow velocity magnitude, and black rods indicate beating directions.

**Movie S5:** Time-evolution of the mucus flow and ciliary-forcing directions, for $\phi = 0.5$ and $\lambda = 4$. The flow is visualized using iso-contours of the flow velocity magnitude, and black rods indicate beating directions.

### Numerical methods

#### Lattice-Boltzmann method

The mucus flow, governed by Eq. (1), is simulated using a lattice-Boltzmann method (Krüger et al., 2017). This method, instead of directly describing the macroscopic flow behavior, is based on a mesoscopic description, namely a statistical description of the microscopic fluid particle dynamics. The particle distribution function $f(\boldsymbol{x}, \boldsymbol{\xi}, t)$ represents the density of fluid particles with velocity $\boldsymbol{\xi}$ at location $\boldsymbol{x}$ and time $t$. Its dynamics is governed by the Boltzmann equation, which in the absence of external forcing writes

$$\frac{\partial f}{\partial t} + \boldsymbol{\xi} \cdot \boldsymbol{\nabla f} = \Gamma(f), \tag{S1}$$

where $\Gamma$ designates the collision operator. At the macroscopic level, the Boltzmann equation is equivalent to the Navier-Stokes equations, and it is thus relevant to describe the present mucus flow.

The discretization of Eq. (S1) in velocity space, physical space and time leads to the lattice-Boltzmann equation. The velocity space is discretized on a set of velocity vectors $\{\boldsymbol{c_l}, l = 0, ..., Q-1\}$, where $Q$ is the number of discrete velocities. In the present work, a $D2Q9$ velocity set is used, in which the velocity space is discretized by nine velocities, namely

$$\boldsymbol{c_l} = \begin{cases} (0,0), & l = 0, \\ c\left(\cos(\frac{\pi(l-1)}{2}), \sin(\frac{\pi(l-1)}{2})\right), & l \in [1,4], \\ \sqrt{2}c\left(\cos(\frac{\pi(2l-9)}{4}), \sin(\frac{\pi(2l-9)}{4})\right), & l \in [5,8], \end{cases} \tag{S2}$$

where $c$ is the lattice speed, and $\boldsymbol{e_x}$ and $\boldsymbol{e_y}$ are unit vectors in the $x$ and $y$ directions. The particle densities at velocities $\{\boldsymbol{c_l}\}$ are represented by the discrete-velocity distribution functions $\{f_l(\boldsymbol{x}, t)\}$, also called particle populations. Time and space are discretized so that particle populations are transported from one node to a neighboring one during one time step, namely $\Delta x/\Delta t = \Delta y/\Delta t = c$. The associated grid is thus Cartesian and uniform. The grid spacing is denoted by $\Delta n = \Delta x = \Delta y$. In the following, all quantities are normalized by $c$ and $\Delta t$, so that $\Delta n = \Delta t = 1$.

Using this normalization, the lattice-Boltzmann equation writes

$$f_l(\boldsymbol{x} + \boldsymbol{c_l}, t+1) - f_l(\boldsymbol{x}, t) = \Gamma_l(\boldsymbol{x}, t), \tag{S3}$$

where $\Gamma_l$ is the discretized collision operator. The left-hand side of (S3) describes the streaming step; the right-hand side is the collision step. Equation (S3) is explicit, and the streaming and collision steps can be treated separately.

The macroscopic flow quantities are moments of the particle populations in the velocity space. In particular, the fluid momentum $\rho_m \boldsymbol{U}$ and density $\rho_m$ write

$$\rho_m \boldsymbol{U} = \sum_{l=0}^{8} f_l \boldsymbol{c_l}, \quad \rho_m = \sum_{l=0}^{8} f_l. \tag{S4}$$

Even though the lattice-Boltzmann method allows small variations of the fluid density, in practice these variations are negligible if the flow velocity remains small, and the fluid can be considered as close to incompressible.

In the present simulations, a Bhatnagar-Gross-Krook (BGK) collision operator is employed, namely

$$\Gamma_l(\boldsymbol{x}, t) = -\frac{1}{\tau}\left(f_l(\boldsymbol{x}, t) - f_l^{(eq)}(\boldsymbol{x}, t)\right), \tag{S5}$$

where $\tau$ is called the relaxation time and determines the kinematic fluid viscosity through $\nu_m = \frac{1}{3}(\tau - 1/2)$, using the present normalization, and $f^{(eq)}$ denotes the equilibrium particle distribution function, expressed as

$$f_l^{(eq)} = w_l \rho_m \left(1 + \frac{\boldsymbol{U}\cdot\boldsymbol{c_l}}{c_s^2} + \frac{(\boldsymbol{U}\cdot\boldsymbol{c_l})^2}{2c_s^4} - \frac{\boldsymbol{U}\cdot\boldsymbol{U}}{2c_s^2}\right). \tag{S6}$$

The weights $\{w_l\}$ are specific to the velocity set. In the present case ($D2Q9$ velocity set), $w_0 = 4/9$, $w_1 = w_2 = w_3 = w_4 = 1/9$ and $w_5 = w_6 = w_7 = w_8 = 1/36$.

#### Inclusion of an external forcing

The effect of the body force $\boldsymbol{f_n}$ is ensured by the external forcing scheme proposed by Guo et al. (2002). The lattice-Boltzmann equation becomes

$$f_l(\boldsymbol{x}+\boldsymbol{c_l}, t+1) - f_l(\boldsymbol{x}, t) = \Gamma_l(\boldsymbol{x}, t) + \left(1 - \frac{1}{2\tau}\right) S_l(\boldsymbol{x}, t), \tag{S7}$$

where $S_l$ is a source term expressed as

$$S_l = w_l \left( \frac{\boldsymbol{c_l} - \boldsymbol{U}}{c_s^2} + \frac{(\boldsymbol{c_l} \cdot \boldsymbol{U}) \cdot \boldsymbol{c_l}}{c_s^4} \right) \cdot \boldsymbol{f_n}. \tag{S8}$$

The flow momentum has to be corrected according to the external forcing, following

$$\rho_m \boldsymbol{U} = \sum_{l=0}^{8} f_l \boldsymbol{c_l} + \frac{1}{2} \boldsymbol{f_n}. \tag{S9}$$

As described in the section *Hydrodynamic model of ciliary-beat organization* of the main manuscript, the forcing $\boldsymbol{f_n}$ is decomposed in two parts, i.e. the ciliary force $\boldsymbol{f_c}$ and the friction force $\boldsymbol{f_v}$. However, the friction depends on the fluid velocity, namely $\boldsymbol{f_v} = -\kappa \boldsymbol{U}$. This term is thus treated implicitly, and the flow momentum is expressed as

$$\rho_m \boldsymbol{U} = \frac{\sum_{l=0}^{8} f_l \boldsymbol{c_l} + \frac{1}{2} \boldsymbol{f_c}}{1 + \kappa/2\rho_m}. \tag{S10}$$

#### Algorithm and numerical parameters

A summary of the implemented algorithm is presented in table 1. During the initialization, the ciliated elements are randomly placed in the computational domain. An example of initialization is shown in Fig. S1(a). A closer view of the hexagonal mesh and underlying fluid nodes is given in Fig. S1(b). The distance between two neighboring nodes, in the $x$ or $y$ direction, is denoted by $\Delta n$; it is set to $\Delta n = D/5$, where $D$ denotes the side length of the hexagonal elements. An hexagonal element thus typically contains 65 fluid nodes. At each time step, the flow velocity is averaged in each ciliated elements, and the angle of the ciliary forcing is updated following Eq. (2). The parameter $\Omega$, which drives the transient re-orientation of the ciliated elements but does not affect the final steady solution in the absence of noise, is set to $\Omega = U_0/D$, unless otherwise stated. A small threshold angle is employed, namely $\theta_0 = 2\Omega\Delta t = 0.004$ radians using the present numerical parameters, so that its influence on model solutions can be neglected. The number of performed time steps varies from one computation to the other, since simulations are systematically continued until a steady solution is reached. This is achieved by checking the statistical convergence of several global quantities as the total fluid kinetic energy, the total flow rate and the space-average instantaneous variation of the ciliary-force orientations.

1. At the beginning of the computation, the fluid velocity is set to zero, and the particle populations are equal to the equilibrium distributions (S6).

2. The hexagonal mesh is initialized, and the ciliated elements are randomly selected according to the prescribed ciliary density. The ciliary forces are initialized with random orientations. The ciliary forces are transferred from the hexagonal mesh to the fluid nodes.

3. The time-marching loop is performed until the required number of time steps is achieved :

3.1. The ciliary forces are updated according to Eq. (2):

3.1.1. On each ciliated cell $i$, the space-averaged flow velocity is computed to determine the local flow orientation angle $\theta_f^i$.

3.1.2. The orientation difference $\Delta\theta^i$ is computed, and the new ciliary force orientation is determined accordingly.

3.1.3. The updated ciliary forces are transferred from the hexagonal mesh to the fluid nodes.

3.2. Collision is performed (S5)

3.3. Streaming is performed (S3).

3.4. Macroscopic quantities are computed (S4,S9), and the friction force $\boldsymbol{f_v}$ is updated.

4. Output of the final solution (flow velocity, ciliary forces, etc).

Table 1: Summary of the present numerical algorithm.

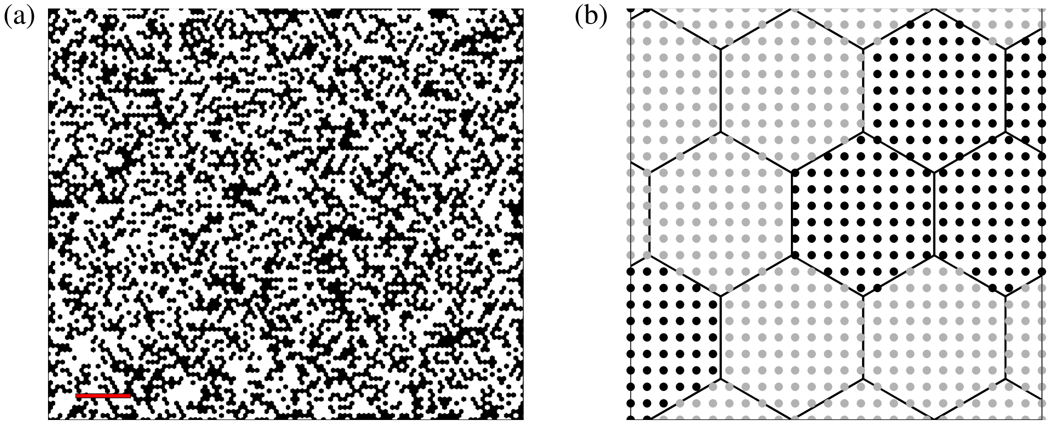


Figure S1: **Spatial discretization employed in the model** (a) Visualization of the numerical domain, for a ciliary density $\phi = 0.4$. The epithelium is discretized using hexagonal elements, representing cells or groups of cells. The ciliated elements, indicated in black, are randomly placed during the initialization of the computations. The scale bar corresponds to $20D$. (b) Closer visualization representing the hexagonal elements and the underlying numerical grid. On ciliated elements, the ciliary forcing is locally imposed on lattice nodes, indicated by black points. On non-ciliated elements (grey points), the ciliary forcing vanishes. The side length of hexagonal elements is equal to $D$.

#### Effect of noise on self-organized regimes

The swirly self-organized regime observed numerically and experimentally appears to be stable in the presence of noise. This is depicted in Figs. S2 and S3. In these simulations, the threshold angle $\theta_0$ is set to 0, so the dynamics of ciliated elements is fully governed by the competition between noise and streamwise alignment (see Eq. (2) in the paper). The magnitude of these mechanisms is controlled by the angular velocity $\Omega$ and the noise intensity $I_n$. Fig. S2 shows the effect of these parameters of the ciliary organization and flow pattern for $\phi = 0.7$ and $\lambda = 1$. In all cases, simulations are initialized using the same steady swirly solution, obtained in the absence of noise. Computations are run until statistical convergence is achieved. It is noted that the self-organized regime remains robust for several sets of parameters, even when the noise magnitude is significantly larger than angular velocity ($I_n = 10\Omega$, Fig. S2(c)). Disorganization of the system is observed for very large relative noise magnitudes, namely ($I_n = 100\Omega$, Fig. S2(a)).

As shown in Fig. S3, a similar behavior is observed for $\phi = 0.45$ and $\lambda = 2$, even though this case is close to the swirly/aligned transition region in the phase diagram (see Fig. 4 in the paper). The system remains almost unaffected for several values of $I_n$ and $\Omega$, and disorganization is only observed for $I_n = 100\Omega$ (Fig. S3(a)).

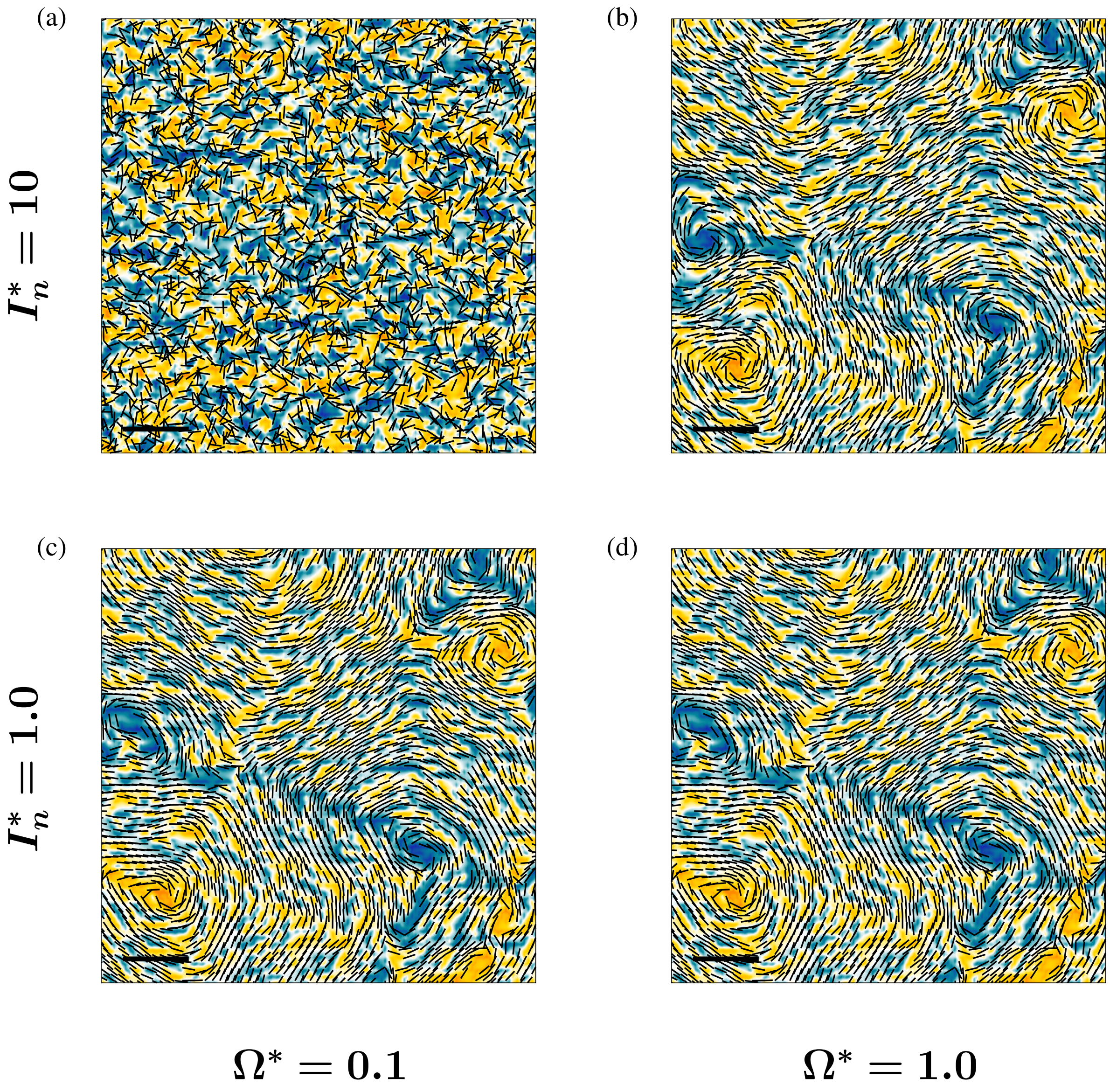


Figure S2: Effect of noise on the swirly regime for $\phi = 0.7$ and $\lambda = 1$. Regimes predicted by the model are visualized using iso-contours of the non-dimensional vorticity ($\omega = [-1, 1]$) and black rods that indicate local beating directions, for various values of $\Omega^* = \Omega D/U_0$ and $I_n^* = I_n D/U_0$. Part of the computational domain is shown and the scale bars correspond to $15D$.

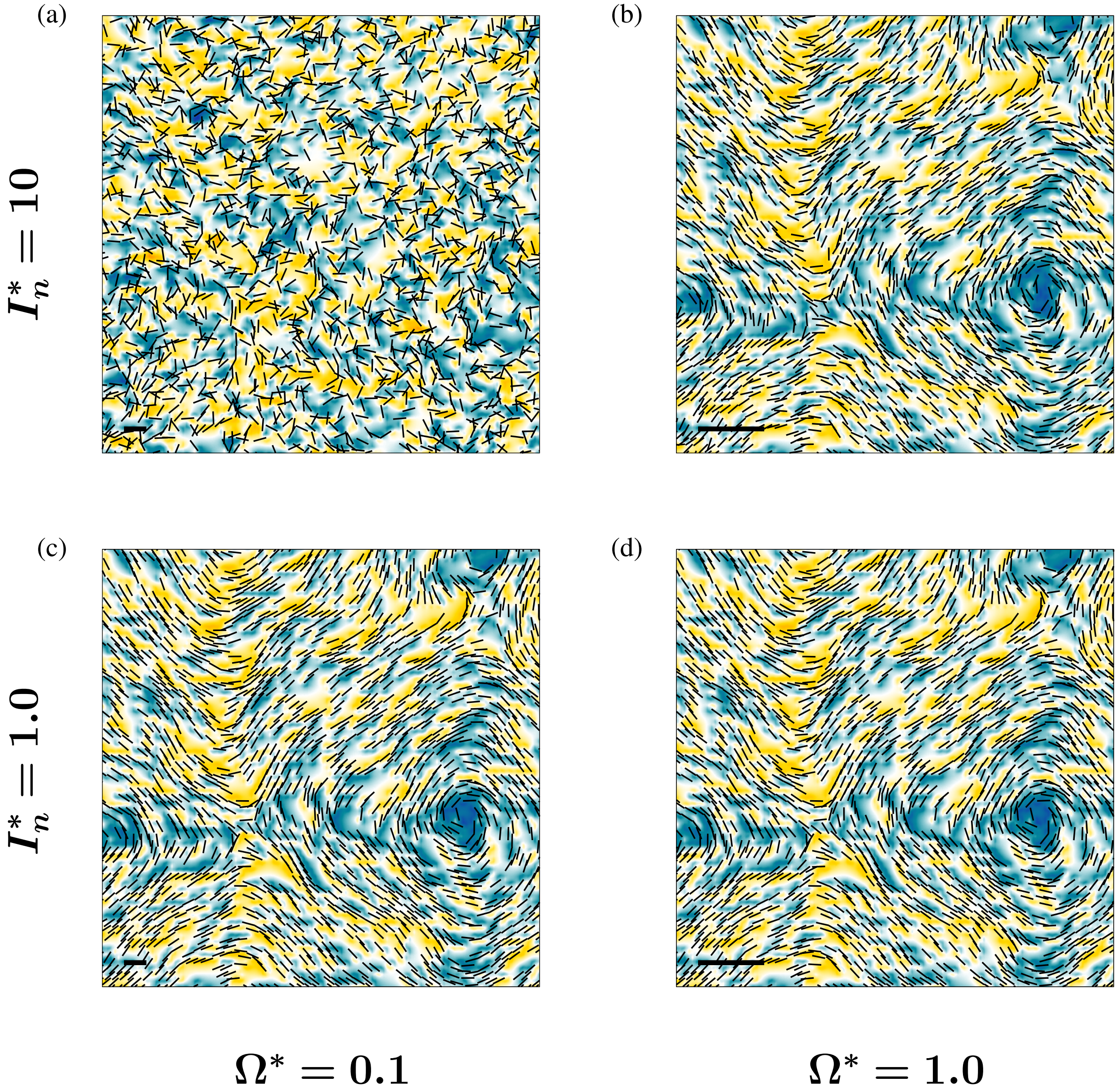


Figure S3: Effect of noise on the swirly regime for $\phi = 0.45$ and $\lambda = 2$. Regimes predicted by the model are visualized using iso-contours of the non-dimensional vorticity ($\omega = [-0.5, 0.5]$) and black rods that indicate local beating directions, for various values of $\Omega^* = \Omega D/U_0$ and $I_n^* = I_n D/U_0$. Part of the computational domain is shown and the scale bars correspond to $15D$.

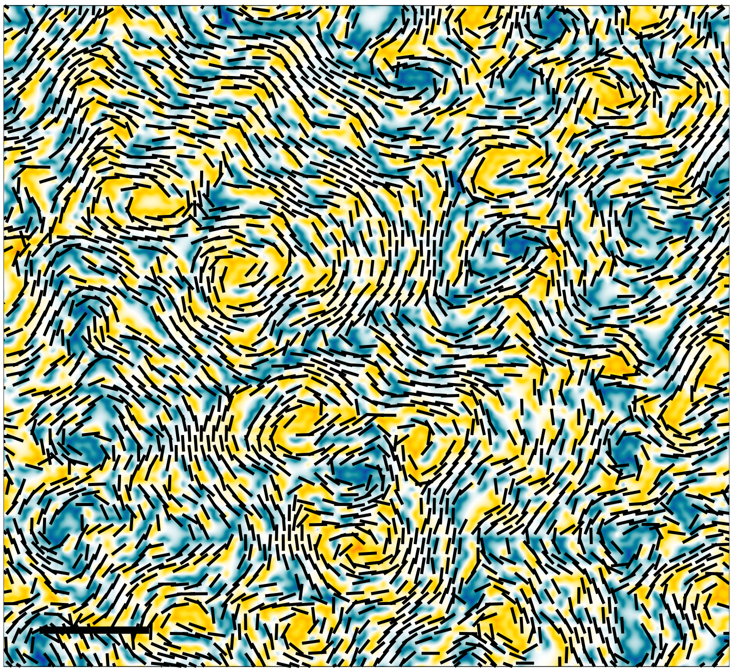

Figure S4: Visualization of the mucus flow and ciliary-beat pattern for $\phi = 0.7$ and $\lambda = 0.5$. This configuration is employed as a model of the system after mucus removal (see Fig. 1(c)), i.e. when $\mu_m = \mu_p$. The flow is visualized using iso-contours of the non-dimensional vorticity ($\omega = [-2, 2]$), and black rods indicate beating directions. Part of the computational domain is shown and the scale bar corresponds to $15D$.

### References

Timm Krüger, Halim Kusumaatmaja, Alexandr Kuzmin, Orest Shardt, Goncalo Silva, and Erlend Magnus Viggen. *The Lattice Boltzmann Method: principles and practice*. Springer, 2017.

Zhaoli Guo, Chuguang Zheng, and Baochang Shi. Discrete lattice effects on the forcing term in the lattice boltzmann method. *Physical Review E*, 65(4):046308, 2002.
